# Thalamus is a common locus of reading, arithmetic, and IQ: Analysis of local intrinsic functional properties

**DOI:** 10.1101/2020.05.05.076232

**Authors:** Maki S. Koyama, Peter J. Molfese, Michael P. Milham, W. Einar Mencl, Kenneth R. Pugh

## Abstract

Neuroimaging studies of basic achievement skills, reading and arithmetic, often control for the effect of IQ to identify unique neural correlates of each skill. This may underestimate possible effects of common factors between achievement and IQ measures on neuroimaging results. Here, we simultaneously examined achievement (reading and arithmetic) and IQ measures in young adults, aiming to identify MRI correlates of their common factors. Resting-state fMRI (rs-fMRI) data were analyzed using two metrics assessing local intrinsic functional properties; regional homogeneity (ReHo) and fractional amplitude low frequency fluctuation (fALFF), measuring local intrinsic functional connectivity and intrinsic functional activity, respectively. ReHo highlighted the thalamus/pulvinar (a subcortical region implied for selective attention) as a common locus for both achievement skills and IQ. More specifically, the higher the ReHo values, the lower the achievement and IQ scores. For fALFF, the left superior parietal lobule, part of the dorsal attention network, was positively associated with reading and IQ. Collectively, our results highlight attention-related regions, particularly the thalamus/pulvinar as a key region related to individual differences in performance on all the three measures. ReHo in the thalamus/pulvinar may serve as a tool to examine brain mechanisms underlying a comorbidity of reading and arithmetic difficulties, which could co-occur with weakness in general intellectual abilities.

**Highlights:**

- Achievement (reading; arithmetic) and IQ measures are simultaneously examined; not controlling for IQ.
- ReHo and fALFF are used to examine local intrinsic functional connectivity and activity, respectively.
- Higher ReHo in thalamus/pulvinar is associated with lower performance on all the three measures.
- Higher fALFF in the left superior parietal lobule is associated with higher word reading and IQ.

## 1. Introduction

Reading and arithmetic are basic achievement skills that influence an individuals’ success at school and beyond. As these domain-specific achievement skills are correlated with general intellectual abilities (indexed by IQ scores) (Gagné & St Père, 2001; Lambert & Spinath, 2018; Susan Dickerson Mayes, Calhoun, Bixler, & Zimmerman, 2009; Peng, Wang, Wang, & Lin, 2019), neuroimaging studies often control for the effect of IQ (i.e., IQ being entered as a covariate of no-interest or being matched between groups) to identify unique neural underpinnings of the achievement skills and their impairments (Ashkenazi, Rosenberg-Lee, Tenison, & Menon, 2012; De Smedt, Holloway, & Ansari, 2011; Eden et al., 2004; Hoeft et al., 2006; Koyama et al., 2011; Pugh et al., 2008; Rosenberg-Lee, Barth, & Menon, 2011). Although this analytical practice has been criticized from logical, statistical, and/or methodological perspectives (Dennis et al., 2009), it remains a topic of debate on whether IQ should be controlled for when studying relationships between brain structures/functions and the achievement skills. This lack of consensus in the literature is evident by the fact that majority of most recent neuroimaging studies of reading and arithmetic have still opted to control for IQ (Ashburn, Flowers, Napoliello, & Eden, 2020; Bulthe et al., 2019; Jolles et al., 2016; Michels, O’Gorman, & Kucian, 2018; Paz-Alonso et al., 2018).

Some prior studies, addressing the role of IQ in predicting the achievement skills and intervention responses, have indicated that IQ is not a direct cause of either academic achievement (Brankaer, Ghesquiere, & De Smedt, 2014; J. M. Fletcher, Francis, Rourke, Shaywitz, & Shaywitz, 1992; Francis, Fletcher, Shaywitz, Shaywitz, & Rourke, 1996; Murayama, Pekrun, Lichtenfeld, & Vom Hofe, 2013) or intervention responses for learning difficulties (Stuebing, Barth, Molfese, Weiss, & Fletcher, 2009; Vellutino, Scanlon, & Lyon, 2000). Furthermore, neuroimaging studies have demonstrated that activations in core regions involved in the achievement skills (e.g., the left temporoparietal junction for reading) are independent of IQ (Hancock, Gabrieli, & Hoeft, 2016; Simos, Rezaie, Papanicolaou, & Fletcher, 2014; Tanaka et al., 2011). These prior findings lead us to think that significant correlations observed between the achievement and IQ tests likely reflect the consequence of both tests measuring common latent factors. Under this circumstance, the use of IQ as a covariate of no-interest could remove some unspecified factors accounting for an achievement skill, and thus potentially producing overcorrected or counterintuitive MRI findings. However, it may be equally misguiding to fail to use IQ as a covariate of interest, which would result in disregarding possible effects of shared factors between achievement and IQ measures on brain activation/connectivity.

Alternatively, both achievement and IQ measures can be simultaneously examined (e.g., an F-test with the two measures of interest) to detect regions where brain signals can be explained by either measure or their combination (Mumford, Poline, & Poldrack, 2015). This approach can answer questions, such as “which regions show significant associations with either measure (e.g., reading or IQ) or both measures”. In particular, the identification of regions common to both measures could help to understand neuromechanisms underlying bidirectional interactions between the achievement and IQ measures. Such bidirectional interactions have been recently appreciated, with mounting evidence from longitudinal studies. For reading and IQ relationships, early reading performance predicts later IQ, and early IQ predicts later reading performance (Chu, vanMarle, & Geary, 2016; Ramsden et al., 2013; Ritchie, Bates, & Plomin, 2015). For arithmetic and IQ relationships, 10-week arithmetic training improves IQ, and 13-week reasoning training improves arithmetic performance (Lowrie, Logan, & Ramful, 2017; Sanchez-Perez et al., 2017). Most evidently, a meta-analysis of longitudinal studies (Peng et al., 2019) has rendered further evidence that intellectual abilities and achievement skills (both reading and mathematics) predict each other even after controlling for initial performance. Crucially, neural substrates underlying such relations, which can be mediated by shared latent factors in the achievement skills and IQ, cannot be delineated by common analytical practices in the literature, that is, IQ being either covaried out (i.e., controlled for) or excluded from analysis.

In the current resting-state functional MRI (rs-fMRI) study, we address this issue by simultaneously examine both achievement (either reading or arithmetic) and IQ measures. Our primary aim is to explore rs-fMRI correlates common to both the achievement and IQ measures in young adults, whose achievement and IQ scores ranged along a continuum from conventionally impaired to superior performance. Specifically, we address our aim in a twofold way; 1) entering two covariates of interest – one for an achievement measure (either reading or arithmetic) and the other for Full-Scale IQ (FSIQ) and 2) entering the first principal component (PC1) – reduced from the three measures (reading, arithmetic, and FSIQ) – as the covariate of interest. The first approach uses F-tests, allowing us to detect regions associated with either measure (i.e., specific) or those associated with the common variance explained by the two measures (Mumford et al., 2015). The second approach using principal component analysis (PCA) allows us to explore common regions (Pugh et al., 2013), reflecting the shared variance among the three measures (i.e., two achievement measures and FSIQ), irrespective of the achievement domains.

When analyzing rs-fMRI data, we focus on two data-driven metrics that index local/regional intrinsic functional properties; the first is voxel-wise regional homogeneity (ReHo; Zang, Jiang, Lu, He, & Tian, 2004), and the second is fractional amplitude of low frequency fluctuations (fALFF; Zou et al., 2008). ReHo, which is calculated with Kendall’s coefficient of concordance (KCC), estimates local or short-distance intrinsic functional connectivity (iFC) between the time-series of a given voxel and its nearest neighboring voxels. Jiang, et al (2015; 2016) have postulated that a higher ReHo value, representing higher synchronization of regional brain activity, indicates higher functional specification in a given region (e.g., the primary visual cortex has the highest ReHo value among regions in the visual ventral pathway). Unlike ReHo, fALFF is a frequency-domain analysis to assess the relative contribution of specific low frequency oscillations to the whole frequency range (Zou et al., 2008). That is, fALFF is a measure of local brain activity, and does not provide any information on functional connectivity. Hence, ReHo and fALFF could be complementary in such that they potentially reveal different brain regions associated with cognitive functions and dysfunctions, although similar results/regions are often reported (Bueno et al., 2019; Hu et al., 2016; Yuan et al., 2013).

Both ReHo and fALFF have successfully detected regions associated with individual differences in cognitive abilities (S. Kuhn, Vanderhasselt, De Raedt, & Gallinat, 2014; Yang et al., 2015), clinical diagnoses/traits (Du, Liu, Hua, & Wu, 2019; Han et al., 2018; Hoexter et al., 2018; Respino et al., 2019; Xu, Zhuo, Qin, Zhu, & Yu, 2015; Xue, Lee, & Guo, 2018), and training/experience effects (Koyama, Ortiz-Mantilla, Roesler, Milham, & Benasich, 2017; Qiu et al., 2019; Salvia et al., 2019; Wu et al., 2019). However, to date, there are only a handful of studies that have applied these metrics to examination of achievement skills, IQ, and/or their relationships. For reading, M. Xu et al. (2015) have examined fALFF, with controlling for IQ, and revealed that positive associations between fALFF in reading-related regions (e.g., the posterior superior temporal gyrus) and semantic reading. For arithmetic, Jolles et al. (2016) compared a group of children with mathematical difficulties (i.e., lower arithmetic abilities) and the IQ-matched control group, the first of which was characterized by higher fALFF in the intraparietal sulcus – a core region associated with number processing (Dehaene, Piazza, Pinel, & Cohen, 2003) and arithmetic (Bugden, Price, McLean, & Ansari, 2012; Dehaene, Molko, Cohen, & Wilson, 2004; Jolles et al., 2016; Menon, 2010). Regarding ReHo, no study has explored its whole-brain patterns associated with either achievement or IQ measures (but see Koyama et al., 2017 using a region of interest analysis).

We opt to use data-driven ReHo and fALFF as the primary metrics, rather than seed-based correlation analysis (SCA) that is the most common way to examine resting-state functional connectivity. This is because ReHo and fALFF require no prior knowledge or hypotheses, unlike SCA that requires the selection of seeds (i.e., regions of interest). Investigators typically select seeds based on previous task-evoked fMRI findings in relevant cognitive domains: for example, Koyama et al. (2011) have employed multiple seeds based on meta-analysis studies of reading-related fMRI findings. This selection of seeds is investigator-specific (e.g., seed location, seed size), making SCA vulnerable to bias. In other words, SCA potentially overlooks brain regions that are not selected by investigators, as well as brain regions that are not typically activated during cognitive tasks of interest. For example, when examining auditory processing and its disorders, SCA would typically use seeds located in the primary auditory cortex based on prior task-evoked fMRI results (Bartel-Friedrich, Broecker, Knoergen, & Koesling, 2010; Talavage, Gonzalez-Castillo, & Scott, 2014); however, Pluta et al. (2014) have highlighted that ReHo in the precuneus, a core region of the default mode network (Buckner, Andrews-Hanna, & Schacter, 2008; Raichle et al., 2001), rather than the auditory cortex, is associated with auditory processing disorders. Based on above-mentioned studies, we hypothesize that the current study, which uses data-driven ReHo/fALFF, could reveal regions outside the networks that had been reported by previous fMRI studies (e.g., task-evoked fMRI, resting-state fMRI using SCA) of the achievement skills and IQ. This possibility can be even enhanced given that we simultaneously examine both achievement and IQ measures (e.g., IQ as a covariate of interest in F-tests), rather than controlling for IQ (e.g., IQ as a covariate of non-interest) – the latter is a long-standing common or standard analytic practice in the neuroimaging literature of reading and arithmetic.

## 2. Materials and Methods

### 2.1. Participants

Seventy-two young adults (28 males; mean age 21±1.9 years, age range = 18-25) were selected from a larger study (N = 159) that primarily aimed to examine neural mechanisms of sequence learning and overnight consolidation in adolescents and young adults. The inclusion criteria used here were as follows; 1) young adults older than 25, 2) the completion of a battery of standardized tests measuring cognitive abilities, at least word reading, arithmetic, and IQ, 3) the completion of two rs-fMRI scans within the same MRI session, 4) English as the first language, and 5) no demonstration of either extreme left-handedness assessed by the Edinburgh Inventory (Oldfield, 1971) (i.e., extreme left-handedness was defined by the score falling between -13 and -24; scores range from -24 to 24, showing the most extreme left- and right-handedness, respectively), and 6) no excessive in-scanner head motion, indexed by mean frame-wise displacement (FD) (Jenkinson, Bannister, Brady, & Smith, 2002) > 0.2mm, in either the first or second rs-fMRI. Demographic characteristics of the participants are given in Table 1. Prior to participation, written informed consent was obtained from all participants in accordance with the guideline provided by Yale University’s Institutional Review Board with the Human Investigation committee. Participants received monetary compensation for their time and effort.

**Table 1.** Demographic and cognitive characteristics of included participants. (N = 72; 28 males) WJ = Woodcock Johnson, WASI = Wechsler Adult Intelligence Scale; FSIQ = Full-Scale IQ, ss = standard score, FD = Framewise Displacement; Rest = Resting-state fMRI

|  | Mean | SD | Range <sup>a</sup> |
| --- | --- | --- | --- |
| Age (years) | 21 | 1.9 | 18.1 – 25.1 |
| Family Income <sup>b</sup> | 2.9 | 1.5 | 1 – 5 |
| Handedness <sup>c</sup> | 12.4 | 1.3 | -12 – 24 |
| WJ Letter Word Identification (ss) | 103.2 | 11.2 | 76 – 124 |
| WJ Calculation (ss) | 102.2 | 15.9 | 68 – 138 |
| WASI FSIQ (ss) | 107.3 | 15.0 | 75 – 145 |
| Mean FD Rest 1 | 0.077 | 0.041 | 0.024 – 0.191 |
| Mean FD Rest 2 | 0.079 | 0.038 | 1.25 – 0.178 |

<sup>a</sup>. “Range” describes the lowest and highest values/scores for each variable in the sample of the current study.
<sup>b</sup>. “Family Income” score ranges from 1 (the lowest) to 5 (the highest).
<sup>c</sup>. “Handedness” score ranges from -24 (the most extreme left-handedness) to 24 (the most extreme right-handedness).

### 2.2. Behavioral measures

One-on-one assessment using standardized measures took place in a quiet room, typically one week prior to the MRI session. Intellectual abilities were assessed using the Wechsler Abbreviated Scale of Intelligence Second Edition (WASI: Wechsler, 1999). In the analysis, we used FSIQ, comprised of both Verbal Intelligence Quotient (VIQ) and Performance Intelligence Quotient (PIQ). The Vocabulary and Similarities subtests are combined to form VIQ, whereas the combination of the Block Design and Matrix Reasoning subtests forms PIQ. The two achievement skills (reading and arithmetic) were assessed using two subtests of the Woodcock-Johnson Tests of Achievement Third Edition (WJ-III) (Woodcock, Mather, & McGrew, 2007); 1) Letter Word Identification (LW) where participants were asked to identify and sound out isolated letters and words from an increasingly difficult vocabulary list and 2) Calculation (Calc) where participants were given a response booklet and asked to complete written mathematical/numerical operations at basic (e.g., addition, division) and higher (e.g., geometric, logarithmic) levels. Standard scores were obtained using age-based norms. Summary statistics for performance on the three measures (LW, Calc, and FSIQ) are provided in Table 1 and Figure 1.

**Figure 1.**
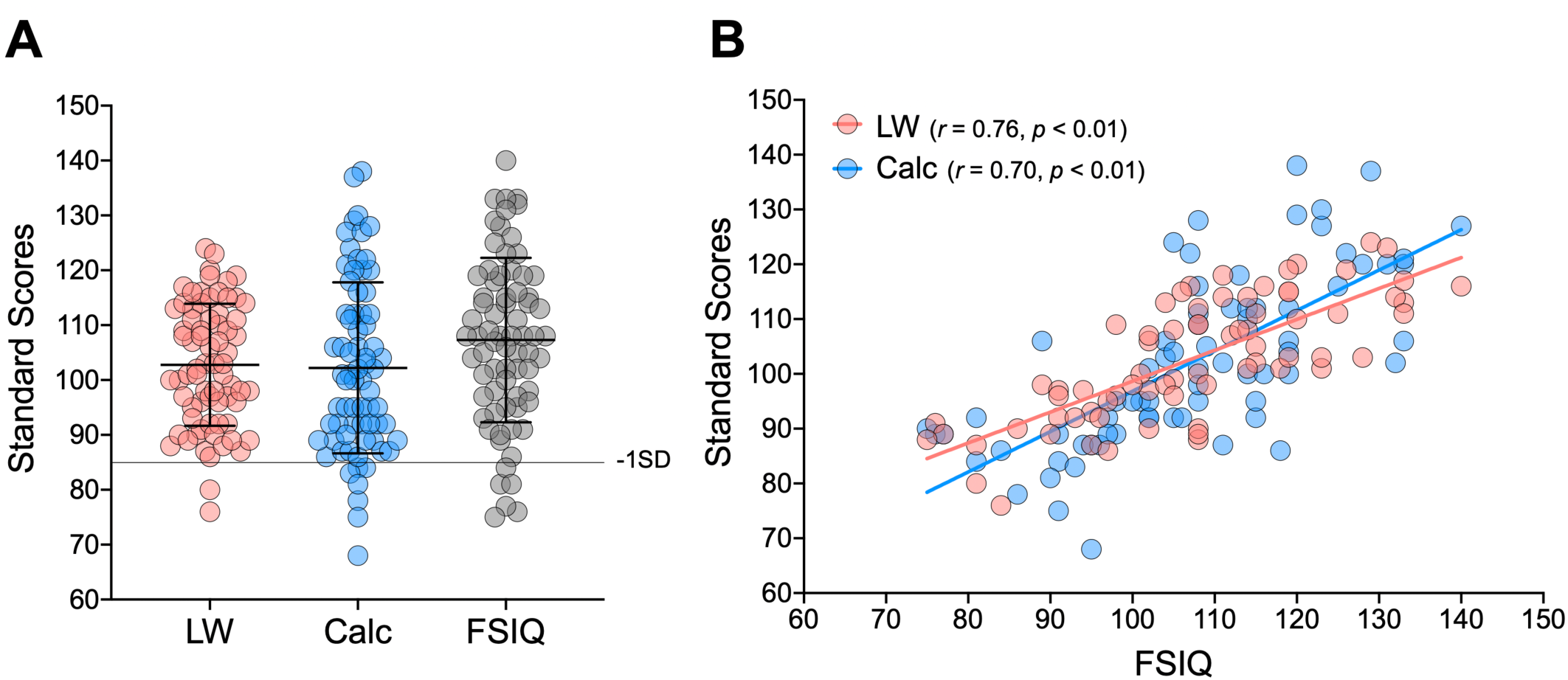
Performance on three cognitive measures (N = 72). **A)** Mean and standard deviation of each measure; Letter Word Identification (LW), Calculation (Calc), and Full-Scale IQ (FSIQ). The horizontal line represents the standard score of 85 (−1SD). **B)** Relationships between FSIQ and the two achievement skills (LW and Calc).

The current analysis included all participants without any cut-off scores on the three measures of interest, mainly for two reasons; 1) impairments with achievement skills, particularly reading, are considered to represent the lower tail of a normal distribution of the abilities (Jack M. Fletcher et al., 1994; Rodgers, 1983; Shaywitz, Escobar, Shaywitz, Fletcher, & Makuch, 1992) (But see Rutter & Yule, 1975; Stevenson, 1988) and 2) the Diagnostic and Statistical Manual of Mental Disorders Fifth Edition (DSM-5: AmericanPsychiatricAssociation, 2013) has de-emphasized specific IQ scores as a diagnostic criterion of Specific Learning Disabilities and Intellectual Disabilities. Accordingly, there were some participants whose standard scores fell below the average range (lower than 85, defined as “weakness’). As shown in Figure 1.A, for FSIQ, none of our participants scored lower than 70 (two standard deviations below the average). This indicates that none of the participants included in the current study had intellectual disability. However, six participants’ scores fell into the 70-84 range, which can be classified as borderline intellectual functioning (Alloway, 2010; Wieland & Zitman, 2016). For reading, two participants demonstrated LW weakness (i.e., standard scores lower than 85), both of whom also scored lower than 85 on FSIQ. For arithmetic, seven participants demonstrated Calc weakness, only one of whom scored lower than 85 on FSIQ. No participant scored lower than 85 on both LW and Calc, that is, no individual in our sample had comorbid weakness in reading and arithmetic.

### 2.3. Principal component analysis (PCA) for LW, Calc, and FSIQ

Consistent with previous findings (Gagné & St Père, 2001; Susan Dickerson Mayes et al., 2009), LW, Calc, and FSIQ were strongly correlated with each other in our sample: LW with FSIQ (*r(70)* = 0.76, *p* < 0.01 in Figure 1.B), Calc with FSIQ (*r(70) =* 0.71, *p* < 0.01 in Figure 1.B), and LW with Calc (*r(70) =* 0.64, *p* < 0.01). Given these strong inter-correlations among the three main measures, we performed PCA, into which standard scores from the three measures were entered. For this analysis, we used the “Scikit-learn 0.19.1” (Python library: Python 3.6.4). Extracted first principal component (PC1) scores were subsequently used in one of statistical models to gain a better understanding of how PC1, the largest shared variance among the three measures, is associated with rs-fMRI metrics.

### 2.4. MRI procedure

Participants received explicit instructions and were placed comfortably in the MRI scanner. To prevent excessive motion during scanning, participants’ head in the head-coil was surrounded by memory foam cushions. To protect their hearing in the MRI scanner, participants wore over-ear headphones in addition to the disposable ear plugs. The MRI session was primarily composed of an 8-min structural MRI scan, two rs-fMRI scans (5 minutes for each scan), and 5-min task-evoked fMRI scans using a serial reaction time task (SRTT). The first rs-fMRI scan was sandwiched between the first and second SRTT scans, while the second rs-fMRI scans took place immediately after the second SRTT scan. During each rs-fMRI scan, participants were instructed to remain still and keep their eyes open. The SRTT employed in the current study was described elsewhere (Hung et al., 2019). Total duration of the MRI session was approximately 45 min.

### 2.5. MRI data acquisition

All MRI data were collected using a Siemens TimTrio 3.0 Tesla scanner located at Yale School of Medicine’s Magnetic Resonance Research Center. Each of the two rs-fMRI scans was comprised of 150 contiguous whole-brain functional volumes acquired using an echo-planar imaging (EPI) sequence (effective TE = 30ms; TR = 2000ms; flip angle = 80°; 32 axial slices; voxel-size = 3.4×3.4×4.0mm; field of view = 220mm). A high-resolution T1-weighted structural image was also acquired using a magnetization prepared gradient echo sequence (MPRAGE, TE = 2.77ms; TR = 2530ms; TI = 1100ms; flip angle = 7°; 176 slices; acquisition voxel size = 1.0×1.0×1.0mm; field of view = 256mm). For each individual, both structural (MPRAGE) and functional (rs-fMRI) data were visually inspected for excessive motion before data preprocessing. Via this visual inspection, 9 participants were excluded from the initial dataset (N = 159).

### 2.6. MRI data preprocessing

Data preprocessing was carried out using the Configurable Pipeline for the Analysis of Connectomes (CPAC version 0.3.9.1 http://fcp-indi.github.io/docs/user/index.html). To allow for stabilization of the magnetic field, the first three volumes within each rs-fMRI dataset were discarded. Our rs-fMRI data preprocessing included the following steps: realignment to the mean EPI image to correct for motion, grand mean-based intensity normalization (all volumes scaled by a factor of 10,000), nuisance regression, spatial normalization, temporal band-pass filtering (0.01–0.1 Hz: this application was only for ReHo because fALFF involves computation of the power across the entire frequency spectrum), and spatial smoothing. For each individual’s rs-fMRI data, mean FD was calculated, and participants who showed “excessive motion” (mean FD > 0.2mm) were excluded from further analyses.

Nuisance regression was performed to control for the effects of head motion and to reduce the influence of signals of no interest. The regression model included linear and quadratic trends, the Friston-24 motion parameters (6 head motion, their values from one time point before, and the 12 corresponding squared items) (Friston, Williams, Howard, Frackowiak, & Turner, 1996), and the signals of five principal components derived from noise regions of interest (e.g., white matter, cerebral spinal fluid) using a component-based noise correction method (CompCor) (Behzadi, Restom, Liau, & Liu, 2007). Spatial normalization included the following steps: (1) anatomical-to-standard registration using Advanced Normalization Tools (ANTs; Avants et al., 2011); (2) functional-to-anatomical registration using FLIRT (Jenkinson et al., 2002) with a 6-degrees of freedom linear transformation, which was further refined using the Boundary-based Registration implemented in FSL (Greve & Fischl, 2009); and (3) functional-to-standard registration by applying the transformation matrices obtained from step (1) and (2) using ANTs. Finally, spatial smoothing was performed, via FSL, using a Gaussian kernel (Full width at half maximum = 8 mm).

### 2.7. ReHo and fALFF

At the individual level, ReHo and fALFF maps were generated for each of the two rs-fMRI datasets obtained, resulting in two ReHo and two fALFF maps for each participant. For ReHo and fALFF, we primarily focused on the first rs-fMRI data and restricted the use of the second rs-fMRI data only to SCA (see “2.10.”) and confirmatory analyses (see “2.11.”). This was because a potential effect of recent task performance would be lesser on the first rs-fMRI than the second rs-fMRI (i.e., participants undertook one SRTT scan prior to the first rs-fMRI, while they had two SRTT scans prior to the second rs-fMRI).

ReHo, an index of local iFC, is defined as KCC for the time series of a given voxel with those of its nearest neighboring voxels (Zang et al., 2004). KCC for each voxel was calculated voxel-wise by applying a cluster size of 26 voxels (faces, edges, and corners) according to the following formula;

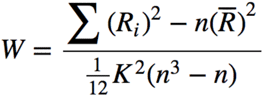

where W was the KCC of given voxels, ranging from 0 to 1, *Ri* was the rank sum of the *ith* time-point; 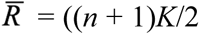 was the mean of the *Ri*, K was the number of time-series within a measured cluster (n = 27; one given voxel plus the others inside the cluster), and n was the number of ranks (corresponding to time-points). For fALFF that measures the intensity of intrinsic functional activity (Zou et al., 2008), we computed the power spectrum at each voxel by transforming the time series to the frequency domain, and then calculated the square root of the amplitude at each frequency (which is proportional to the power at that frequency). Finally, we divided the sum of the amplitude across the low frequencies (0.01 to 0.1 Hz) by the sum of the amplitudes across the entire frequency range. For each participant, ReHo and fALFF were computed in the native space, registered in the MNI space, and then smoothed. At the group level, we used FSL’s FEAT to perform whole-brain voxel-wise general linear models, with a study-specific mask that included voxels (in MNI space) present in at least 90% of participants. Whole-brain correction for multiple comparisons was performed using Gaussian Random Field Theory (*Z* > 3.1; cluster significance of *p* < 0.05).

### 2.8. Group models

We performed the following three group models to examine both ReHo and fALFF; 1) LW and FISQ (i.e., standard scores) entered as covariates of interest into an F-test, 2) Calc and FSIQ (i.e., standard scores) entered as covariates of interest into an F-test, and 3) PC1 entered as the covariate of interest. In each group model, we additionally included covariates of non-interest; age, sex, mean FD, and whole-brain mean of each rs-fMRI metric (Yan, Milham 2013). For both of the first and second models, we used FSIQ although the literature has shown that VIQ correlates more strongly with reading than does PIQ, whereas PIQ correlates more strongly with arithmetic skills than does VIQ (Ashkenazi, Rosenberg-Lee, Metcalfe, Swigart, & Menon, 2013; Strauss, Sherman, & Spreen, 2006). The main rationale for using FSIQ was that the correlation between FSIQ and LW (*r(70)* = 0.76, *p* < 0.01) was not significantly different from the correlation between VIQ and LW (*r(70)* = 0.77, *p* < 0.01) (*Z* = 0.29, *p* = 0.38) (Lenhard & Lenhard, 2014). Similarly, the correlation between FSIQ and Calc (*r(70)* = 0.71, *p* < 0.01) was not significantly different from the correlation between PIQ and Calc (*r(70) =* 0.67, *p* < 0.01) (*Z* = 0.93, *p* = 0.17). For the third model, we performed PCA to detect a single factor (i.e., PC1) underlying common variation among the three measures that were significantly correlated with each other. Results of PCA showed that the PC1 accounted for 81% of the total variance (Note that principal component 2 accounted only for 12%). PC1 scores extracted for each individual were significantly (*p* < 0.01) correlated with LW (*r(70)* = 0.83), Calc (*r(70)* = 0.88), and FISQ (*r(70)* = 0.94), confirming that the PC1 is relevant to the achievement and intellectual abilities in the current sample. Resultant clusters were labelled/defined based on the Harvard-Oxford Cortical and Subcortical Structural Atlases, as well as the Colin27 Subcortical Atlas modified from the one described in Chakravarty, Bertrand, Hodge, Sadikot, and Collins (2006).

### 2.9. Brain-behavior relationships

We extracted the mean ReHo/fALFF value across all voxels within each significant F-test result/cluster from each participant, and then calculated *R*-squared and *p* values that represent brain-behavior relationships. As F-test results are non-directional (i.e., positive or negative), we made post-hoc visualization by plotting the mean ReHo/fALFF values extracted from each significant result/cluster as a function of the achievement skills and FSIQ. In addition, ReHo and fALFF values extracted from each significant cluster identified by the model with PC1 (i.e., positively and/or negatively associated with PC1 scores) were plotted as a function of the achievement skills and FSIQ.

### 2.10. Seed-based correlation analysis (SCA)

We selected seed regions based on significant results from the whole-brain ReHo/fALFF analyses with the first rs-fMRI data. Importantly, we used the second rs-fMRI data for post-hoc SCA in order to avoid “double dipping” in our fMRI analysis (Kriegeskorte, Simmons, Bellgowan, & Baker, 2009). Our SCA aimed to explore whether and how global or longer-distance iFC of given seeds (i.e., significant results from local rs-fMRI metrics) would be associated with LW, Calc, and/or FSIQ. At the individual level, the average time series across the voxels within each seed was extracted and correlated with all voxels within the group-specific mask, using Pearson’s correlation. Correlation values were transformed to Fisher *Z* scores to provide a whole-brain iFC map of each seed for each participant. At the group level, we employed the above-mentioned three models. The resultant iFC maps were corrected for multiple comparisons using Gaussian Random Field Theory (*Z* > 3.1; cluster significance: *p* < 0.05).

### 2.11. Confirmatory region of interest (ROI) analysis using the second rs-fMRI data

We applied ROI analysis to the second rs-fMRI data collected from the same sample, and then examined if results (i.e., brain-behavior relationships) from the whole-brain analysis using the first rs-fMRI data were significant in the second rs-fMRI data. This confirmatory ROI analysis aimed to test intra-individual reliability of brain-behavior relationships. For this purpose, we extracted the mean values of the second rs-fMRI ReHo/fALFF from each of significant clusters detected in the analyses using the first rs-MRI data. Subsequently, we calculated *R*-squared and *p* values that represent brain-behavior relationships in the second rs-fMRI data.

## 3. Results

### 3.1. ReHo results from the first rs-fMRI

Table 2 and Figure 2 summarize significant ReHo results from the three models. Both F-tests (i.e., the first and second models) highlighted the thalamus with the peak voxels located in the left pulvinar (the “LW-Thalamus” cluster for the F-test with LW & FSIQ; the “Calc-Thalamus” cluster for the F-test with Calc & FSIQ). Additionally, the PC1 scores were negatively associated with the thalamus (the “PC1-Thalamus” cluster).

**Table 2.** Significant ReHo results from the three models. LW = Letter Word Identification, Calc = Calculation, FSIQ = Full-Scale IQ, PC1 = Principal Component 1

| Contrast | Brain Region | Peak MNI Coordinates | # of Voxels |
| --- | --- | --- | --- |
| LW & FSIQ (F-test) | Thalamus/Left Pulvinar | -12 -27 -3 | 171 |
| Calc & FSIQ (F-test) | Thalamus/Left Pulvinar | -12 -27 -6 | 144 |
| PC1 (Negative correlation) | Thalamus/Left Pulvinar | -12 -27 -3 | 168 |

**Figure 2.**
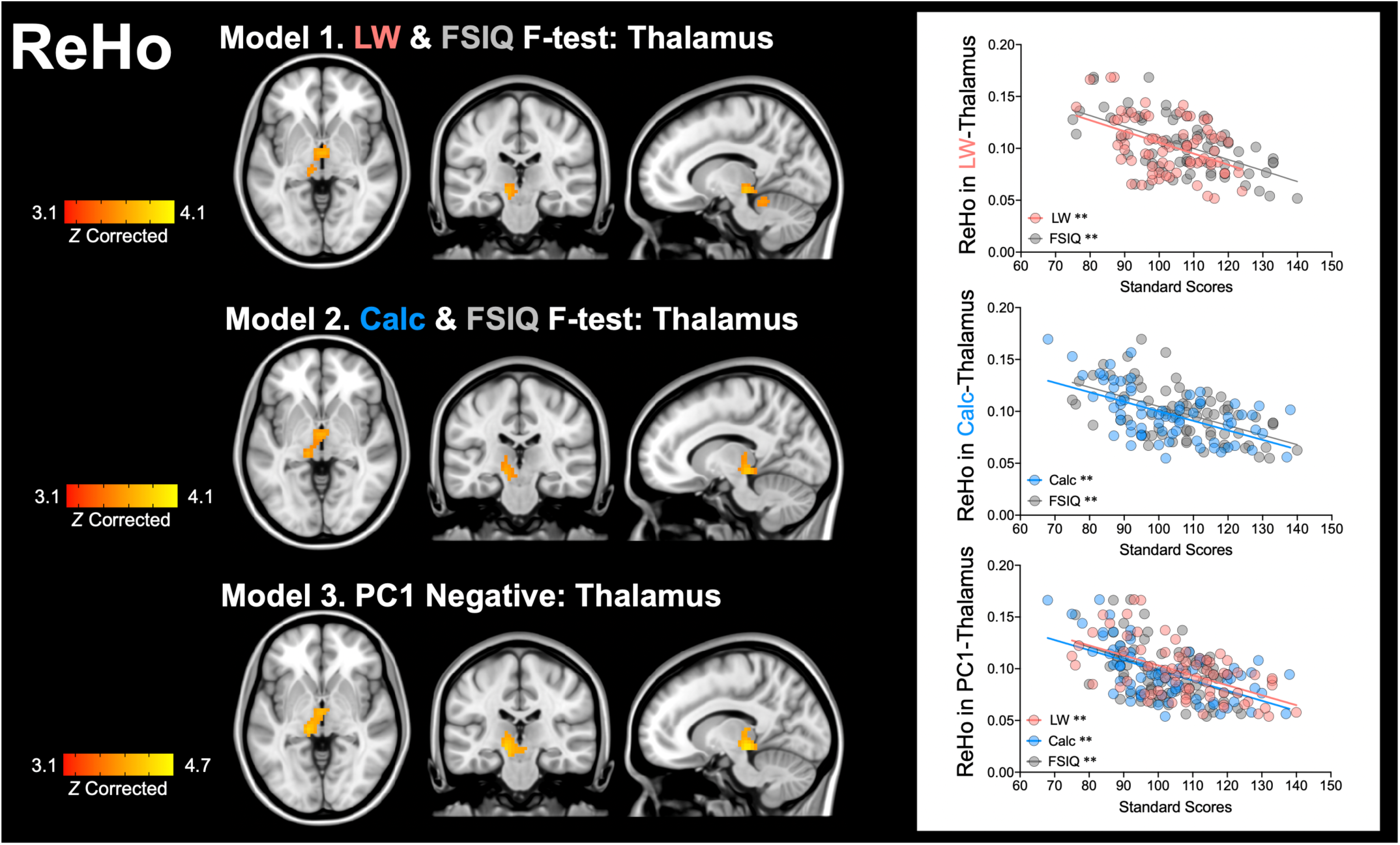
Significant ReHo results from the three models. Results from all the models, mapped on the MNI coordinates (x = -12, y = -27, z = -3), highlight the thalamus, centered in the left pulvinar. Model 1 performs an F-test with two covariates of interests – Letter Word Identification (LW) and Full-Scale IQ (FSIQ). Model 2 performs an F-test with Calculation (Calc) and FSIQ. In Model 3, the first principal component (PC1) among three measures (LW, Calc, and FSIQ) is entered as the covariate of interest. In scatter plots on the right, mean ReHo values extracted from the respective thalamus cluster are plotted as a function of LW, Calc and FSIQ. Images are displayed according to neurological convention (left is left). ** *p* < 0.001

### 3.2. ReHo-behavior relationships

We plotted the mean ReHo values (i.e., the first rs-fMRI data) extracted from each cluster as a function of the corresponding cognitive measures (e.g., ReHo values in the LW-Thalamus cluster as a function of LW and FSIQ), as shown in the upper and middle scatter plots in Figure 2. First, the ReHo values from the LW-Thalamus cluster were negatively associated with LW (*R*^*2*^ = 0.20, *p* < 0.001) and FSIQ (*R*^*2*^ = 0.35, *p* < 0.001): these two associations, with one variable in common (i.e., ReHo in the LW-Thalamus), were significantly different (*Z* = 2.1, *p* < 0.05); more strongly associated with FSIQ than LW. Second, the ReHo values from the Calc-Thalamus cluster were negatively associated with Calc (*R*^*2*^ = 0.32, *p* < 0.001) and FSIQ (*R*^*2*^ = 0.30, *p* < 0.001); these two associations, with one variable in common (i.e., ReHo in the Calc-Thalamus), were not significantly different (*Z* = 0.27, *p* = 0.78). In addition, we visualized relationships for the PC1 result by plotting the ReHo values from the PC1-Thalamus cluster as a function of each measure (LW, Calc, and FSIQ). There were significant negative associations for all three measures; LW (*R*^*2*^ = 0.21, *p* < 0.001), Calc (*R*^*2*^ = 0.33, *p* < 0.001) and FSIQ (*R*^*2*^ = 0.30, *p* < 0.001) (The bottom scatter plot in Figure 2). These three associations, with one variable in common (i.e., ReHo in the PC1-Thalamus), were not significantly different between one another (*Z* = 1.14, *p* = 0.15 for LW and FSIQ; *Z* = 0.41, *p* = 0.68 for Cal and FSIQ; *Z* = 1.43, *p* = 0.15 for LW and Calc).

### 3.3. Common thalamus and ReHo results from the second rs-fMRI

As ReHo results from all the three models highlighted the thalamus, we identified the overlap among the three thalamus clusters and then created the “Common-Thalamus” cluster, which was located dominantly in the left hemisphere (the image on the left in Figure 3). In the first rs-fMRI data, ReHo values from the Common-Thalamus cluster was significantly associated with LW (*R*^*2*^ = 0.13, *p* < 0.01), Calc (*R*^*2*^ = 0.21, *p* < 0.001), and FSIQ (*R*^*2*^ = 0.22, *p* < 0.001). These three associations, with one variable in common (i.e., ReHo in the Common-Thalamus), were not significantly different from one another (*Z* = 1.47, *p* = 0.15 for LW and FSIQ; *Z* = 0.25, *p* = 0.80 for Cal and FSIQ; *Z* = 1.09, *p* = 0.27 for LW and Calc). This pattern (i.e., no significant differences in the associations) was seen at the thalamus cluster derived from the PC1 – the largest shared variance among the three measures.

**Figure 3.**
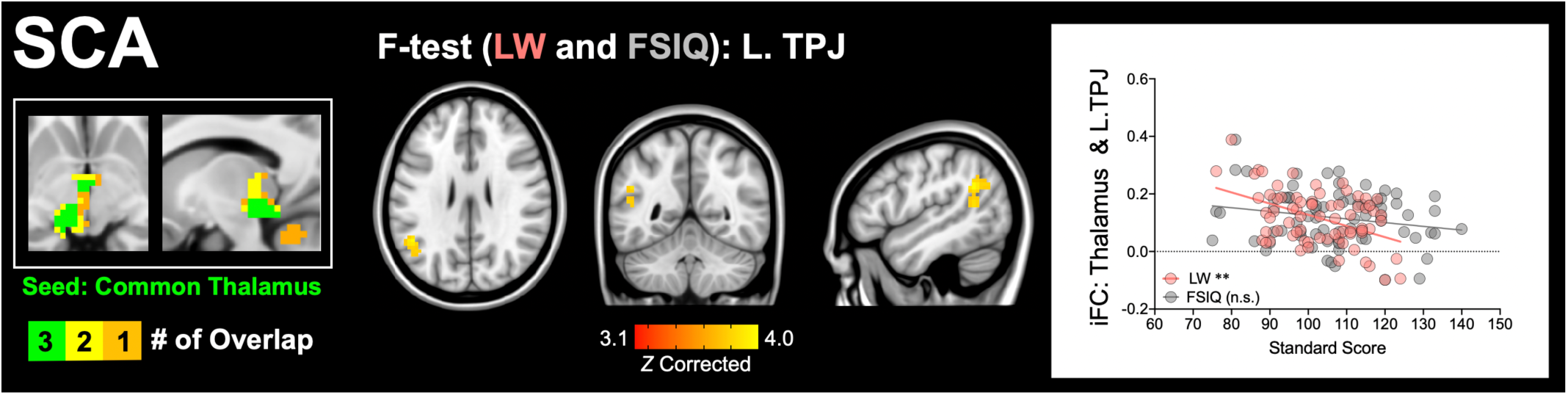
Common-Thalamus cluster and significant seed-based correlation analysis (SCA) results. The overlap among the three thalamus clusters is highlighted in green (the left image). The result from an F-test with Letter Word Identification (LW) and Full-Scale IQ (FSIQ), mapped on the MNI coordinates (x=-46, y=-52, z=26), highlights intrinsic functional connectivity (iFC) between the Common-Thalamus cluster and left temporoparietal junction (L.TPJ). In the scatter plot on the right, iFC values extracted from the thalamus-L.TPJ connectivity are plotted as a function of LW and FSIQ. Images are displayed according to neurological convention (left is left). ** *p* < 0.001, n.s. = not significant

To test reliability of our thalamus results, we performed confirmatory ROI analyses, using the second rs-fMRI data, for four thalamus clusters identified by the whole-brain analysis using the first rs-fMRI data: 1) LW-Thalamus, 2) Calc-Thalamus, 3) PC1-Thalamus, and 4) Common-Thalamus. Results showed significant (*p* < 0.001) ReHo-behavior relationships in the second rs-fMRI data: 1) the LW-Thalamus cluster with LW (*R*^*2*^ = 0.19) and FSIQ (*R*^*2*^ = 0.24), 2) the Calc-Thalamus cluster with Calc (*R*^*2*^ = 0.20) and FSIQ (*R*^*2*^ = 0.29), 3) the PC1-Thalamus cluster with LW (*R*^*2*^ = 0.19), Calc (*R*^*2*^ = 0.24), and FSIQ (*R*^*2*^ = 0.30), and 4) the Common-Thalamus cluster with LW (*R*^*2*^ = 0.16), Calc (*R*^*2*^ = 0.20) and FSIQ (*R*^*2*^ = 0.21). Taken together, the results from both the first and second rs-fMRI datasets suggest that higher ReHo in the thalamus is reliably associated with lower LW, Calc, and FSIQ.

### 3.4. fALFF results from the first and second rs-fMRI

As shown in Figure 4, the F-test with LW and FSIQ highlighted the left superior parietal lobule (L.SPL: peak voxel at x = -42, y = -54, z = 51 in MNI). The fALFF values extracted from the L.SPL cluster exhibited a significant positive association with LW (*R*^*2*^ = 0.25, *p* < 0.01) and FSIQ (*R*^*2*^ = 0.06, *p* < 0.05); these two associations, with one variable in common (i.e., fALFF in L.SPL), were significantly different (*Z* = 3.55, *p* < 0.001); more strongly associated with LW than FSIQ. These results remained significant in the second rs-fMRI (*R*^*2*^ = 0.24, *p* < 0.001 for LW; *R*^*2*^ = 0.09, *p* < 0.05 for FSIQ). There was no significant result from either the F-test with Calc and FSIQ or PC1.

**Figure 4.**
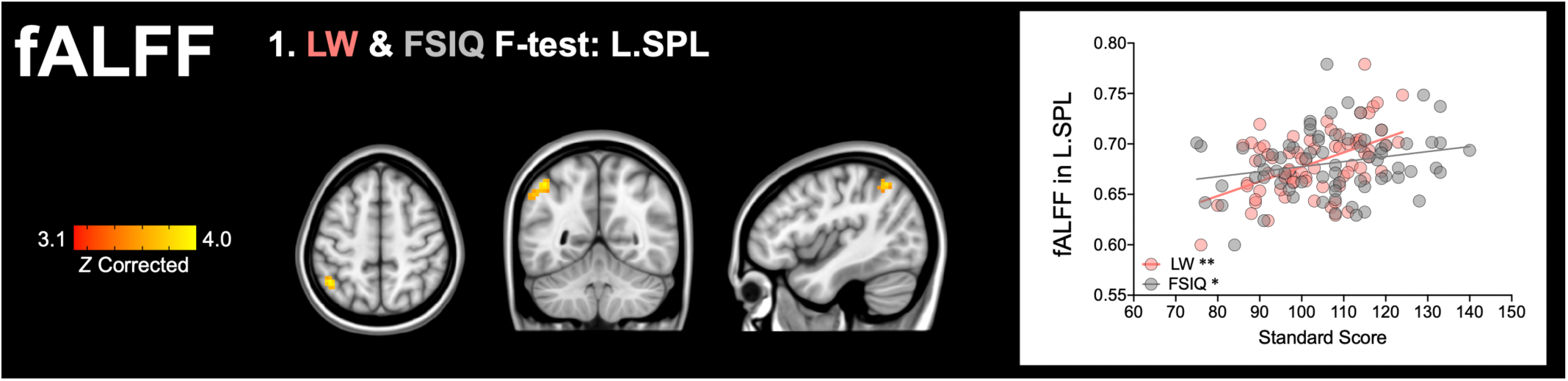
Significant fALFF result. The result from an F-test with Letter Word Identification (LW) and Full-Scale IQ (FSIQ), mapped on the MNI coordinates (x = -42, y = -54, z = 51), highlights the left superior parietal lobule (L.SPL). In the scatter plot on the right, fALFF values extracted from the L.SPL cluster are plotted as a function of LW and FSIQ. Images are displayed according to neurological convention (left is left). ** *p* < 0.001, * *p* < 0.05

### 3.5. SCA with the common thalamus cluster

Across the three models (i.e., two F-tests and PC1), ReHo results highlighted the thalamus, but there was no such a common region identified by the fALFF analysis. Therefore, we used only the Common-Thalamus cluster as the seeds in post-hoc SCA. As shown in Figure 3, the F-test with LW and FSIQ highlighted a significant cluster with the peak voxel at x = -46, y = -52, z = 26 (MNI), which can be considered as centered in left angular gyrus based on the Harvard-Oxford Cortical Structural Atlas (i.e., the maximum probability of 0.25). However, we labelled it as the left temporoparietal junction (L.TPJ) because it 1) included both parietal and superior temporal regions, and 2) spatially corresponded to a subregion of the left temporoparietal junction identified by rs-fMRI parcellation analysis (Igelstrom, Webb, & Graziano, 2015). Note that the closest similarity between the given two regions/clusters was defined by the smallest Euclidean distance (d = 10.4) calculated using the voxel peak MNI coordinates. The post-hoc analysis and visualization revealed a significant negative relationship between the thalamus-L.TPJ iFC and LW (*R*^*2*^ = 0.21, *p* < 0.001), that is, the higher the iFC, the worse the reading. This negative iFC-behavior relationship was not significant for FSIQ (*R*^*2*^ = 0.04, *p* = 0.08); the two associations, with one variable in common (i.e., iFC), were significantly different (*Z* = 3.27, *p* < 0.001).

We found no significant SCA results from either the model with F-test with Calc/FSIQ or the PC1 model. This absence of significant SCA results was somewhat surprising given that ReHo in the Common-Thalamus cluster was significantly associated with all the three measures (LW, Calc and FSIQ). Thus, we performed post-hoc analyses to examine whether and the degree of which iFC values in the thalamus-L.TPJ connectivity would be associated with Calc and FSIQ scores. We found a significant connectivity-behavior association only with Calc (*R*^*2*^ = 0.10, *p* < 0.01) but not with FSIQ (*R*^*2*^ = 0.04, *p* = 0.08).

## 4. Discussion

The current study simultaneously examined the achievement (reading, arithmetic) and IQ measures in young adults, aiming to identify MRI correlates of their common factors. For this aim, we used F-tests, into each of which an achievement measure and FSIQ were entered, as well as investigating the effect of PC1 among the three measures (reading, arithmetic, and FSIQ). The main finding, which was reliable across these analytic models, is that lower ReHo in the thalamus (the peak voxel in the left pulvinar) was associated with higher performance on each of the three measures. This indicates that the thalamus represents a neural correlate of the shared factor among reading, arithmetic, and FSIQ. This ReHo result centered to the thalamus is partially consistent with our hypothesis that our approach could identify regions outside functional brain networks that had been frequently reported by prior fMRI studies of reading arithmetic, and IQ. In the literature, the thalamus may not be a core region implied for reading, arithmetic, or IQ, but the neuroscientific community has increasingly recognized potentially important role of the thalamus in learning (T. Rose & Bonhoeffer, 2018) and language processes (Klostermann, Krugel, & Ehlen, 2013; Llano, 2013), particularly reading (Achal, Hoeft, & Bray, 2016; Diaz, Hintz, Kiebel, & von Kriegstein, 2012; Gaab, Gabrieli, Deutsch, Tallal, & Temple, 2007; Pugh et al., 2013; Stein, 2018a). We discuss more details and implications of these findings, as well as other findings, in the following sections.

### 4.1. ReHo in thalamus/pulvinar

ReHo results from all the three models suggest that higher ReHo in the thalamus is associated with lower performance on word reading, arithmetic, and FSIQ measures. This pattern of negative brain-behavior relationships is consistent with previous rs-fMRI studies, which have reported that higher ReHo in subcortical regions (and higher mean ReHo in the entire brain) is associated with worse outcomes, such as lower abilities (Dajani & Uddin, 2016; S. Kuhn et al., 2014; Zhao et al., 2019). In particular, Zhao et al. (2019) have demonstrated that higher ReHo in the thalamus was associated with severer symptoms in schizophrenic patients. This finding, together with our ReHo finding, indicates that higher ReHo in the thalamus may index lower functions, possibly across different behavioral domains. Although ReHo is considered to index local functional coupling and reflects the degree of local functional specialization (Jiang et al., 2015; Jiang & Zuo, 2016), the underlying mechanism of ReHo-behavior relationships remains largely unknown. In the next paragraphs, we will debate a possible scenario that could explain the observed negative ReHo-behavior relationships (i.e., the higher the ReHo, the worse the performance).

The brain adaptively reconfigures or changes its functional connectivity between globally distributed regions/networks in response to task demands (Cohen & D’Esposito, 2016; Cole, Bassett, Power, Braver, & Petersen, 2014; Hearne, Cocchi, Zalesky, & Mattingley, 2017; Krienen, Yeo, & Buckner, 2014). Yet, functional connectivity patterns detected at rest correspond well with co-activation patterns during tasks; only ∼40% of connections change (Cole et al., 2014; Krienen et al., 2014; S. M. Smith et al., 2009). Such relatively small but reliable changes in functional network reconfiguration between rest and task seem to contribute to individual differences in behavior. For example, Schulz and Cole (2016) have demonstrated that smaller differences in functional connectivity between rest and task, which reflect more efficient reconfiguration (i.e., less energy required for the change/shift), are associated with higher performance on a variety of cognitive tasks (e.g., language, reasoning). This can be interpreted in such that, in high-performing individuals, their functional connectivity patterns during rest are “preconfigured” similar to those during tasks, and that greater similarity may facilitate more efficient or less energy-costing reconfiguration in the presence of task demand.

Such reconfiguration efficiency or optimized preconfiguration at rest is likely to vary significantly across individuals in the thalamus that exhibits the most notable differences between functional connectivity at rest (i.e., temporal correlations) and on task (i.e., co-activation patterns) (Di, Gohel, Kim, & Biswal, 2013). Specifically, the thalamus is more globally connected during task performance, but is more locally connected at rest (Di et al., 2013). Based on these observations and the aforementioned finding (Schultz & Cole, 2016), we consider that higher ReHo in the thalamus at rest may reflect less optimized/efficient local preconfiguration (i.e., more energy-costing reconfiguration in the presence of a task), which in turn is associated with lower performance. In other words, individuals whose local intrinsic functional connectivity patterns of the thalamus are more similar between rest and tasks are likely to perform better on the achievement and IQ measures. Further studies are needed to test this possibility by analyzing fMRI data collected during both rest and relevant tasks (e.g., word reading) in the same individuals.

A question arising here is the common role of the thalamus among reading, arithmetic, and IQ measures. Recent efforts in the rs-fMRI field have highlighted the thalamus as an integrative hub that is connected with multiple cortical networks (Garrett, Epp, Perry, & Lindenberger, 2018; Greene et al., 2019; Hwang, Bertolero, Liu, & D’Esposito, 2017; Seitzman et al., 2020). As such, meta-analysis of a large database of fMRI activations has revealed that the thalamus is involved in multiple cognitive functions (Hwang 2017). Activation of the thalamus has been sometimes (but not always) reported during tasks that require higher cognitive processing, such as language (Crosson, 2013; Klostermann et al., 2013; Llano, 2013), reading (Gaab et al., 2007; Houde, Rossi, Lubin, & Joliot, 2010; Maisog, Einbinder, Flowers, Turkeltaub, & Eden, 2008; Martin, Schurz, Kronbichler, & Richlan, 2015; Pugh et al., 2008; Pugh et al., 2013), arithmetic (Arsalidou & Taylor, 2011), and intelligence/reasoning (Fangmeier, Knauff, Ruff, & Sloutsky, 2006). Although the thalamus is not typically considered as a core region for reading, arithmetic or intellectual abilities, thalamic abnormalities (i.e., geniculate nuclei) have been noted in individuals with reading disorders (Diaz et al., 2012; Stein, 2018b), as well as those with ADHD (Ivanov et al., 2010; X. Li et al., 2012) that is often comorbid with learning difficulties in reading and/or arithmetic (S. D. Mayes, Calhoun, & Crowell, 2000; Wadsworth, DeFries, Willcutt, Pennington, & Olson, 2015).

Our ReHo results emphasize the left posterior thalamus, corresponding to the location of the left pulivnar (the peak voxels from the three models). Thus, our discussion here focuses on the role of pulvinar and its possible contribution to reading, arithmetic, and intellectual skills. The pulvinar has been most intensively examined in its relation with selective attention (Fischer & Whitney, 2012; Halassa & Kastner, 2017; Kastner et al., 2004; Smith, Cotton, Bruno, & Moutsiana, 2009) – the ability to filter/suppress distracting information and enhance relevant information for goal-oriented task performance. Selective attention is not only a survival instinct across all species (Krauzlis, Bogadhi, Herman, & Bollimunta, 2018) but also implicated in higher cognitive abilities unique to humans, including phonological processing (Yoncheva, Maurer, Zevin, & McCandliss, 2014), reading (Commodari, 2017), and intelligence (Kirk, Gray, Ellis, Taffe, & Cornish, 2016; Unsworth, Fukuda, Awh, & Vogel, 2014). The importance of selective attention in reading, arithmetic, and language is extensively reviewed and discussed elsewhere (Stevens & Bavelier, 2012). In particular, close relationships between selective attention and reading (e.g., visual word recognition) has been well-documented (Commodari, 2017; Graboi & Lisman, 2003). For example, during visual word recognition, selective attention enables a reader to compare the visual/orthographic representation of a written word to his/her mental lexicon (i.e., a list of many relevant and irrelevant words stored in memory) until the match is found. Without selective attention, book and web pages would be merely full of visual clutter. Importantly, selective attention occurs in the auditory (Pugh et al., 1996; von Kriegstein, Eger, Kleinschmidt, & Giraud, 2003) and semantic domains (Rogalsky & Hickok, 2009), as well as integrated domains (i.e., audiovisual semantic) (Y. Li et al., 2016). As such, selective attention in the auditory (or auditory semantic) domain, rather than the visual domain, may be more relevant to FSIQ given that some subtests forming FSIQ are orally presented (e.g., the Vocabulary and Similarities subtests).

Returning to neurobiological mechanisms of selective attention, which requires the involvement of multiple functions in a coordinate fashion and efficient cortico-cortical communications (Yantis, 2008), the pulvinar has been suggested to play a role in selective attention by regulating cortical synchrony (Saalmann & Kastner, 2011; Saalmann, Pinsk, Wang, Li, & Kastner, 2012; Zhou, Schafer, & Desimone, 2016). Such cortical regulation of the pulvinar is most likely to achieved by its extensive interconnection with cortical regions (Greene et al., 2019; Hwang et al., 2017; Seitzman et al., 2020). Although the pulvinar has been prominently examined and discussed in the visual domain (Kaas & Lyon, 2007; Zhou et al., 2016), emerging evidence has suggested its involvement in auditory attention (e.g., speech segmentation) (Dietrich, Hertrich, & Ackermann, 2013, 2015; Erb, Henry, Eisner, & Obleser, 2012). Hence, selective attention, regulated by the pulvinar, is likely to be a common latent factor associated with the reading, arithmetic, and IQ measures. Here, we postulate that higher ReHo in the thalamus (centered in the left pulvinar), reflecting less optimized local preconfiguration at rest, may be associated with lower selective attention across multiple domains, which may exert negative impact on the achievement and IQ performance. This view could, in turn, contribute to the debate as to why some (but not all) individuals with impaired selective attention, such as ADHD (Brodeur & Pond, 2001; Mueller, Hong, Shepard, & Moore, 2017), perform poorly on the achievement and IQ tests (Frazier, Demaree, & Youngstrom, 2004). Validation of this hypothesis deserves further investigation, particularly given that learning difficulties with reading/arithmetic and attention deficits are often understudied or underestimated in people with borderline intellectual functioning (Al-Khudairi, Perera, Solomou, & Courtenay, 2019; Baglio et al., 2014; Di Blasi, Buono, Cantagallo, Di Filippo, & Zoccolotti, 2019; Jansen, De Lange, & Van der Molen, 2013; E. Rose, Bramham, Young, Paliokostas, & Xenitidis, 2009); such individuals are not often granted eligibility to access specialized healthcare or social/educational services they might need (Martinez & Quirk, 2009).

It is worth mentioning that ReHo-behavior relationships may be true only in adults given the reported difference in the amplitude of whole-brain ReHo between children and adults (i.e., lower in adults than children) (Dajani & Uddin, 2016), which is possibly due to pruning of local connections (Jiang et al., 2015) .

Furthermore, developmental changes have been observed in thalamocortical iFC (Fair et al., 2010) and its relationships with reading (e.g., negatively and positively associated with reading in adults and children, respectively) (Koyama et al., 2011); these prior findings implicate possible child-adult differences in patterns of ReHo in the thalamus. Furthermore, given that the thalamus is vulnerable following preterm birth (Ball et al., 2012; Ball et al., 2013; Smyser et al., 2010; Toulmin et al., 2015; Volpe, 2009) and that smaller thalamic volumes at term-equivalent age predict childhood neurodevelopmental deficits (in reading, arithmetic, and IQ) in preterm-born individuals (Loh et al., 2017), early alterations in thalamic functional connectivity may also indicate higher risk for later neurodevelopmental disorders, including learning and intellectual disabilities. It is of great interest to explore the nexus between the development of the thalamus/pulvinar and neurodevelopmental outcomes; this effort would potentially add further evidence for brain mechanisms underlying the comorbidity in reading and arithmetic difficulties (Skeide, Evans, Mei, Abrams, & Menon, 2018; Willcutt et al., 2013), which could co-occur with cognitive weakness in domain-general skills, such as attention, working memory, and speed processing (Gathercole et al., 2016; Willcutt et al., 2013) – these cognitive components are often embedded in IQ tests.

### 4.2. fALFF in L.SPL

The result of the F-test with LW and FSIQ measures suggests that higher fALFF in L.SPL characterizes better word reading and IQ. This positive relationship between local intrinsic activity and reading/IQ is in line with prior task-evoked activation seen in L.SPL during reading (Martin et al., 2015; Meyler, Keller, Cherkassky, Gabrieli, & Just, 2008; Peyrin, Demonet, N’Guyen-Morel, Le Bas, & Valdois, 2011; Reilhac, Peyrin, Demonet, & Valdois, 2013; Vigneau, Jobard, Mazoyer, & Tzourio-Mazoyer, 2005) and intelligence/reasoning tasks (Fangmeier et al., 2006; Goel & Dolan, 2001; Jung & Haier, 2007; Wendelken, 2014). This finding might provide an additional support for evidence that local intrinsic activity, represented by low-frequency oscillations during rs-fMRI, predicts local task activation and relevant behaviors (Kalcher et al., 2013; Mennes et al., 2011). It is well-documented that SPL is a highly heterogeneous region in its patterns of functional connectivity and activation (Caspers & Zilles, 2018; Mars et al., 2011; Scheperjans, Eickhoff, et al., 2008; Wang et al., 2015). Wang et al. (2015) have used a connectivity-based parcellation scheme and detected multiple subregions of SPL, one of which is similar to the cytoarchitectonically defined area 7A (Scheperjans, Eickhoff, et al., 2008; Scheperjans, Hermann, et al., 2008) and appears to correspond to the L.SPL cluster identified in this study. This SPL subregion, particularly in the left hemisphere, has been found to be associated with visuospatial attention, reading, and reasoning (Wang et al., 2015), indicating that the involvement of L.SPL in reading and reasoning may be mediated by visuospatial attention.

SPL is part of the dorsal attention network (Corbetta & Shulman, 2002; Shomstein, 2012; Vossel, Geng, & Fink, 2014), which has a significant overlap with task-positive networks (Dosenbach, Fair, Cohen, Schlaggar, & Petersen, 2008; Dosenbach et al., 2007; Dosenbach et al., 2006; Fox, Corbetta, Snyder, Vincent, & Raichle, 2006), including the frontoparietal network. More recently, Dixon et al. (2018) have differentiated the frontoparietal network in two subsystems, one of which coactivates and is more strongly connected with the dorsal attention network, including SPL. This attention-related subsystem is contrasted to another subsystem that is prominently connected with distributed components of the default mode network. These authors have further explored associations between each subsystem and different cognitive processes, finding that “reading” exhibits a strong positive association with the attention-related subsystem (Dixon et al., 2018). Although some processes (e.g., “working memory”), which can be components measured by IQ tests, are also strongly associated with the attention-related system, these associations are not as strong as the “reading” association. These association patterns are consistent with our finding that the correlation between fALFF in L.SPL and FSIQ were significant but weaker than that for reading. Taken together, our fALFF result suggests that higher local intrinsic activity in L.SPL indexes higher reading and IQ, most likely due to the L.SPL’s involvement in visuospatial attentional control (Corbetta & Shulman, 2002; Wang et al., 2015). This assumption needs to be tested in future research in which aspects of attention (e.g., sustained attention, selective attention) are linked to fALFF in L.SPL (and other attention-related regions).

To date, neurobiological mechanisms of fALFF may be the least understood among rs-fMRI metrics, but recent efforts have revealed that fALFF and ReHo exhibit high correlations with each other (Nugent, Martinez, D’Alfonso, Zarate, & Theodore, 2015; Yuan et al., 2013) and with cerebral blood flow (Z. Li, Zhu, Childress, Detre, & Wang, 2012) at the whole-brain level. Nevertheless, these two local rs-fMRI metrics do not always yield similar or consistent brain regions (and their relationships with behavior in the same populations) (Y. Xu et al., 2015; Yang et al., 2015). This indicates that fALFF and ReHo can provide complementary information about local intrinsic properties, and this is the case in the present study in such that ReHo and fALFF revealed different regions associated with reading (i.e., the thalamus for ReHo and L.SPL for fALFF), as well as different brain-behavior relationships (i.e., positive for ReHo in the thalamus but negative for fALFF in L.SPL).

### 4.3. SCA with the common-thalamus cluster

We found that iFC between the Common-Thalamus cluster and L.TPJ exhibited a significant negative association with LW. That is, individuals with weaker positive correlations (and stronger negative correlations) between these subcortical-cortical regions tended to perform better on the reading measure. This SCA finding is largely consistent with prior rs-fMRI work showing that connections between subcortical (i.e., caudate, thalamus) and left temporoparietal/parietal regions are negatively associated with reading skills in adults (Achal et al., 2016; Koyama et al., 2011). L.TPJ has been widely reported in reading research, particularly due to its hypoactivation in individuals with reading difficulties (Maisog et al., 2008; Martin et al., 2015; Richlan, Kronbichler, & Wimmer, 2009). However, L.TPJ is not specific to reading but is involved in the number of different cognitive functions (Bzdok et al., 2016; Igelstrom & Graziano, 2017), including social reasoning (Samson, et al. 2004) that relies on the default mode network (Buckner, et al. 2008; Li, et al. 2014).

The term “TPJ” is an abstract label that has been commonly used in the neuroimaging literature. Notably, there are differences in labeling the location of TPJ across studies (e.g., different labels describe the same or similar location). As such, Church, Coalson, Lugar, Petersen, and Schlaggar (2008) have reported that, during reading tasks, adult readers show no activation in the left angular gyrus, which is within close proximity of (or overlap with) our L.TPJ cluster: the Euclidean distance = 10.8 based on the peak voxel MNI coordinates. This task-evoked fMRI finding implies that our SCA result of the thalamus-L.TPJ iFC cannot be explained by coactivation patterns (Liu, Zhang, Chang, & Duyn, 2018; S. M. Smith et al., 2009) during reading. Instead, it is possibly linked with the default mode network considering that the L.TPJ cluster identified in this study resembles most closely one of the L.TPJ subdivisions reported in Igelstrom et al. (2015), which is strongly connected and coactivated with the default mode network. Given that stronger negative correlations between task-positive and default mode networks during rest can reflect higher cognitive efficiency (Kelly, Uddin, Biswal, Castellanos, & Milham, 2008), stronger negative iFC (and weaker positive iFC) between the thalamus (i.e., a task-positive region) and our L.TPJ cluster (i.e., a L.TPJ subdivision as part of the default mode network) may reflect improved cognitive efficiency; this is likely to facilitate automaticity, which is an important component of skilled reading (M. R. Kuhn, Schwanenflugel, & Meisinger, 2010; Logan, 1978; Wolf, 2018).

Finally, it is somewhat surprising that the F-test with Calc and FSIQ yielded no significant result in SCA, despite a fact that the thalamus cluster used in SCA was commonly associated with the three measures in the ReHo analysis. However, post-hoc analyses revealed that iFC values extracted from the thalamus-L.TPJ connectivity were significantly associated with arithmetic scores (but not FSIQ). This may imply that there was a possible association between the thalamus-L.TPJ iFC and arithmetic, but it failed to survive correction for multiple comparisons when examining at the whole-brain level (i.e., fMRI data comprising numerous voxels). This result from the post-hoc analysis encourages a future research to have a larger sample size and perform a whole-brain analysis, testing if the thalamus-L.TPJ connectivity could be commonly associated with reading and arithmetic. In addition, given that coactivation patterns of the pulvinar are different according to task types (Barron, Eickhoff, Clos, & Fox, 2015), it will be of great interest to investigate global/long-distance functional connectivity of the thalamus/pulvinar during different tasks, including reading, arithmetic, and intellectual tasks.

## 5. Limitations

There are several limitations in the current study. The most evident was a lack of task-evoked fMRI data in the domain of reading, arithmetic, and IQ/reasoning, which restricted direct comparisons of ReHo-behavior relationships during rest and task. Our primary ReHo finding sits in the thalamus, which is the region exhibiting the most dynamic differences in functional network configuration between rest and task (i.e., more globally connected during task than at rest). Thus, examination of both intrinsic and task-evoked functional connectivity of the thalamus in the same sample could enable us to illustrate comprehensive connectivity-behavior relationships (e.g., a possibility that global/long-distance task-evoked connectivity of the thalamus is positively associated with cognitive abilities). Similarly, although we attributed the ReHo result in the thalamus to the common involvement of selective attention in the achievement and IQ measure, we administered no selective attention skills to be linked to the ReHo result in the thalamus. Given that results from both ReHo and fALFF (i.e., L.SPL in the dorsal attention network) highlight regions involved in attention, which is a prerequisite of learning (Merkley, Matusz, & Scerif, 2018; Reynolds & Besner, 2006; Shaywitz & Shaywitz, 2008), future research studies should consider the measurement of different aspects of attention (e.g., sustained attention, selective attention) and relate them to brain’s functional profiles that characterize reading, arithmetic, and/or IQ. Finally, our results should be interpreted with caution when studying the developing brain (i.e., children); activation and connectivity patterns of the regions identified in the current study are known to be developmentally sensitive and differentially associated with children and adults when reading (Church et al., 2008; Koyama et al., 2011) and arithmetic (Rivera, Reiss, Eckert, & Menon, 2005) abilities are examined.

## 6. Conclusions

We simultaneously examine both achievement (reading, arithmetic) and IQ measures, using rs-fMRI metrics that characterize local intrinsic functional properties. The main finding highlights that ReHo (i.e., local functional connectivity) in the thalamus, particularly the left pulvinar implied in selective attention, is a common neural correlate or convergence site for cognitive variation in reading, arithmetic, and IQ measures. Specifically, the higher the ReHo, the lower the performance on all the three measures. Considering that the thalamus is more locally connected at rest than during tasks, negative ReHo-behavior relationships indicate that higher ReHo in the thalamus at rest may reflect less optimized/efficient local preconfiguration (i.e., more energy-costing reconfiguration in the presence of a task), which in turn is associated with lower performance on each dimension. In addition, the fALFF result suggests that higher local intrinsic functional activity in the left superior parietal lobule (in the dorsal attention network) characterizes better reading and IQ performance. To summarize, our findings provide additional support to claims that attentional components are critical for achievement skills and IQ. In particular, the ReHo finding that the thalamus is a common locus for the three measures could provide a new perspective on brain mechanisms underlying a type of comorbidity between reading and arithmetic difficulties, which could co-occur with weakness in general intellectual abilities.

## Acknowledgements

This study is supported by Eunice Kennedy Shriver National Institute of Child Health and Development (NICHD) R01HD065794 to Haskins Laboratories (Kenneth R. Pugh, PI). We thank Airey Lau, Alexis Lee, Bonnie Buis, Morgan Bontrager, Jocelyn Springfield, and Annie Stutzman for behavioral assessment, recruitment, data collection, and data management. Additional thanks to Jocelyn Springfield for proofreading.

